# Disabling PSGL-1 abrogates immune suppression and resistance to PD-1 blockade in pancreatic cancer

**DOI:** 10.1101/2025.05.22.655365

**Authors:** Jennifer L. Hope, Yijuan Zhang, Hannah A. F. Hetrick, Evelyn S. Sanchez-Hernandez, Beatrice Silvestri, Brianna J. Smith, Sanmati H. Nakil, Sreeja Roy, Michelle Lin, Ashley B. Palete, Swetha Maganti, Lily Ling, Dennis C. Otero, Katelyn T. Byrne, Gabriele Romano, Yu Xin Wang, Cosimo Commisso, Linda M. Bradley

**Affiliations:** Cancer Metabolism and Microenvironment Program, NCI-Designated Cancer Center, Sanford Burnham Prebys, La Jolla, CA, USA; Department of Microbiology and Immunology, College of Medicine, Drexel University, Philadelphia, PA, USA; Immune Cell Regulation and Targeting Program, NCI-Designated Comprehensive Cancer Center, Sidney Kimmel Cancer Consortium, Thomas Jefferson University and Drexel University, Philadelphia, PA, USA; Center for Cardiovascular and Muscular Diseases, Sanford Burnham Prebys, La Jolla, CA USA; Center for Data Sciences, Sanford Burnham Prebys, La Jolla, CA, USA; Cell, Developmental, and Cancer Biology, OHSU Knight Cancer Institute, School of Medicine, Oregon Health and Science University, Portland, OR, USA; Department of Pharmacology and Physiology, College of Medicine, Drexel University, Philadelphia, PA, USA

## Abstract

Pancreatic ductal adenocarcinoma (PDAC) is a lethal cancer for which there is a critical need to identify novel therapeutic targets. Herein we define PSGL-1 as a checkpoint inhibitor using a syngeneic orthotopic model of PDAC. As with PDAC patients, CD8^+^ T cells within murine PDAC tumors expressed high levels of PSGL-1. PSGL-1^-/-^ mice displayed striking T cell-dependent control of primary tumors and lung metastases. Extensive spatial remodeling within PDAC tumors occurred in PSGL-1^-/-^ mice with a dramatic loss of proliferating tumor cells and an increase in CD8^+^ T cell engagement of antigen-presenting cells. The prominent CD8^+^ T cell infiltrates included subsets of pre-exhausted T cells retaining hallmarks of stemness and multifunctional effector capacity. These changes enabled a near complete response of PDAC to therapeutic PD-1 blockade. Our findings identify PSGL-1 as a key regulator of anti-tumor immunity in PDAC, highlighting its potential as a therapeutic target to limit CD8^+^ T cell exhaustion and enhance immunotherapy response.

**Summary:** Hope *et al* describe a pivotal function of PSGL-1 in CD8^+^ T cell responses to pancreatic ductal adenocarcinoma. Genetic deletion of PSGL-1 elicits tumor control by increasing T cell infiltration and maintaining functional subsets, thereby promoting sensitivity to PD-1 blockade.

## Introduction

Pancreatic ductal adenocarcinoma (PDAC) is an aggressive, lethal cancer that has the lowest 5-year survival rate of all adult cancers and is estimated to become the second leading cause of cancer-related deaths in the United States by 2030 (Rahib et al., 2021). PDAC is refractory to all current treatments including immune checkpoint blockade (ICB) targeting PD-1 (Morrison et al., 2018). The poor prognosis of PDAC patients underscores the urgent need to better understand the fundamental mechanisms regulating PDAC to develop innovative therapeutics. Cytotoxic CD8^+^ T cells that destroy tumor cells are critical for protection against cancer, including PDAC. The PDAC tumor microenvironment (TME) is notoriously immunologically cold with infiltrates of functionally exhausted CD8^+^ T cells (Saka et al., 2020) and suppressive myeloid cells (Bear et al., 2020). Thus, it is crucial to identify treatments that overcome T cell exhaustion and neutralize immunosuppressive factors within the PDAC TME.

Exhausted CD8^+^ T cells (T_EX_) develop in response to chronic antigen stimulation. T_EX_ cells progressively upregulate multiple inhibitory receptors (IRs) that include PD-1, TIM-3, and LAG-3, and concomitantly lose effector T cell (T_EFF_) cytotoxicity, cytokine production, and proliferative potential to ultimately become terminally differentiated (Saeidi et al., 2018; Virgin et al., 2009). T_EX_ are now recognized as a distinct CD8^+^ T cell lineage whose differentiation is controlled by unique transcriptional and epigenetic programs that are distinct from those governing effector and memory cells (Belk et al., 2022). T_EX_ are highly heterogenous, consisting of subsets of cells at different stages which differentiate from PD-1 expressing progenitor exhausted cells (T_PEX_) with the capacity for T_EFF_ activity (Chen et al., 2019; Hudson et al., 2019; Utzschneider et al., 2016). These cells are distinguished by expression of the TCF-1 transcription factor (Kallies et al., 2020). Clinically, elevated frequencies of T_PEX_ correlate with cancer patient responses to PD-1 ICB in other solid tumors (Sade-Feldman et al., 2018; Thommen et al., 2018). However, induction of T_PEX_ responses by anti-PD-1 can cause rapid proliferation and differentiation of terminal T_EX_ (Blackburn et al., 2008; Verma et al., 2019). Thus, a fundamental question is whether T_PEX_ and their functional progeny can be generated and maintained to promote, as well as sustain patient responses to ICB therapy. At present, there are no clinically relevant strategies to elicit T_PEX_ in sufficient levels to impact responses to immunotherapy. In this study, we endeavor to fill this critical gap by targeting PSGL-1 (P-selectin glycoprotein-1) in PDAC.

PSGL-1 is primarily expressed by hematopoietic cells and is constitutive on the surface of immune cells including macrophages, dendritic cells, and neutrophils as well as naïve T cells where its baseline expression is rapidly increased upon T cell receptor (TCR) stimulation. We previously identified PSGL-1 as a T cell-intrinsic checkpoint inhibitor that acts upstream of PD-1 to constrain TCR signaling, thereby driving upregulation of multiple IRs and terminal differentiation during chronic antigen stimulation independently of its function in leukocyte migration (Hope et al., 2023; Tinoco et al., 2016). We showed that PSGL-1 engagement in the context of TCR signaling drives T_EX_ development from both mouse and human T_EFF_ *in vitro*, underscoring its integral connection to immune inhibitory pathways and relevance to human anti-tumor responses.

Herein, we show that the gene encoding PSGL-1, *SELPLG*, is highly expressed in tumor infiltrating immune cells in human PDAC and correlates with poor patient survival. To investigate the basis for this association, we used a syngeneic orthotopic PDAC model that recapitulates multiple aspects of human PDAC, including an immune suppressive environment, and resistance to PD-1 ICB (Evans et al., 2016). We show that PSGL-1^-/-^ mice display significant T cell-dependent control of PDAC tumor growth and reduced lung metastases. Compared to tumors from wild type (WT) mice, we found extensive spatial remodeling of the PDAC TME in PSGL-1^-/-^ mice wherein increased frequencies of CD8^+^ T cells were associated with antigen-presenting dendritic cells (APC) and the loss of proliferating tumor cells. CD8^+^ T cells were enriched in TCF-1^+^ T_PEX_, and in CX3CR1^+^ multifunctional T_EFF_. With this shift away from terminal exhaustion, we show that therapeutic treatment of PSGL-1^-/-^ mice with PD-1 ICB resulted in eradication of PDAC tumors but was ineffective in WT control mice. Thus, disabling PSGL-1 was sufficient to remodel the PDAC landscape, to enable significant changes in T_EX_ differentiation that were accompanied by tumor growth control, and to support responses to PD-1 ICB. As such, targeting PSGL-1 may be a novel immunotherapy for ameliorating resistance to PD-1 in PDAC and potentially other refractory solid tumors.

## Results and Discussion

### PSGL-1 expression by PDAC infiltrating immune cells in patients and mice

We evaluated PSGL-1 gene (*SELPLG*) transcript expression in human PDAC tumors and normal tissue using TCGA (The Cancer Genome Atlas), Genotype-Tissue Expression (GTEx), and Gene Expression Profiling Interactive Analysis (GEPIA) databases. *SELPLG* curated datasets reflecting high *SELPLG* expression in PDAC tumors (n=179) compared to normal pancreas (n=171). Kaplan-Meier plots reveal a correlation of high *SELPLG* expression with shorter patient survival (**Fig. 1A**, left panel). We next queried whether high vs low PSGL-1 expression in these tumors were indicative of survival and found that elevated expression was associated with poorer survival (HR = 1.74, p = 0.014) (**Fig. 1A**, right panel). Since PSGL-1 is expressed by most hematopoietic cell, we analyzed publicly available single cell sequencing (scRNA-seq) data from 6 treatment naïve PDAC patients, generating a UMAP based on the 2000 most variable genes identifying 16 Louvain clusters (**Fig. 1B**). *SELPLG* (PSGL-1) expression profiling (blue) demonstrated its restriction to immune infiltrates of which T cells, macrophages, and neutrophils predominated (**Fig. 1B)**. These findings confirm that PSGL-1 is expressed only by immune cells in PDAC infiltrates and that PSGL-1 is a novel patient-relevant immunotherapeutic target.

**Figure 1:**
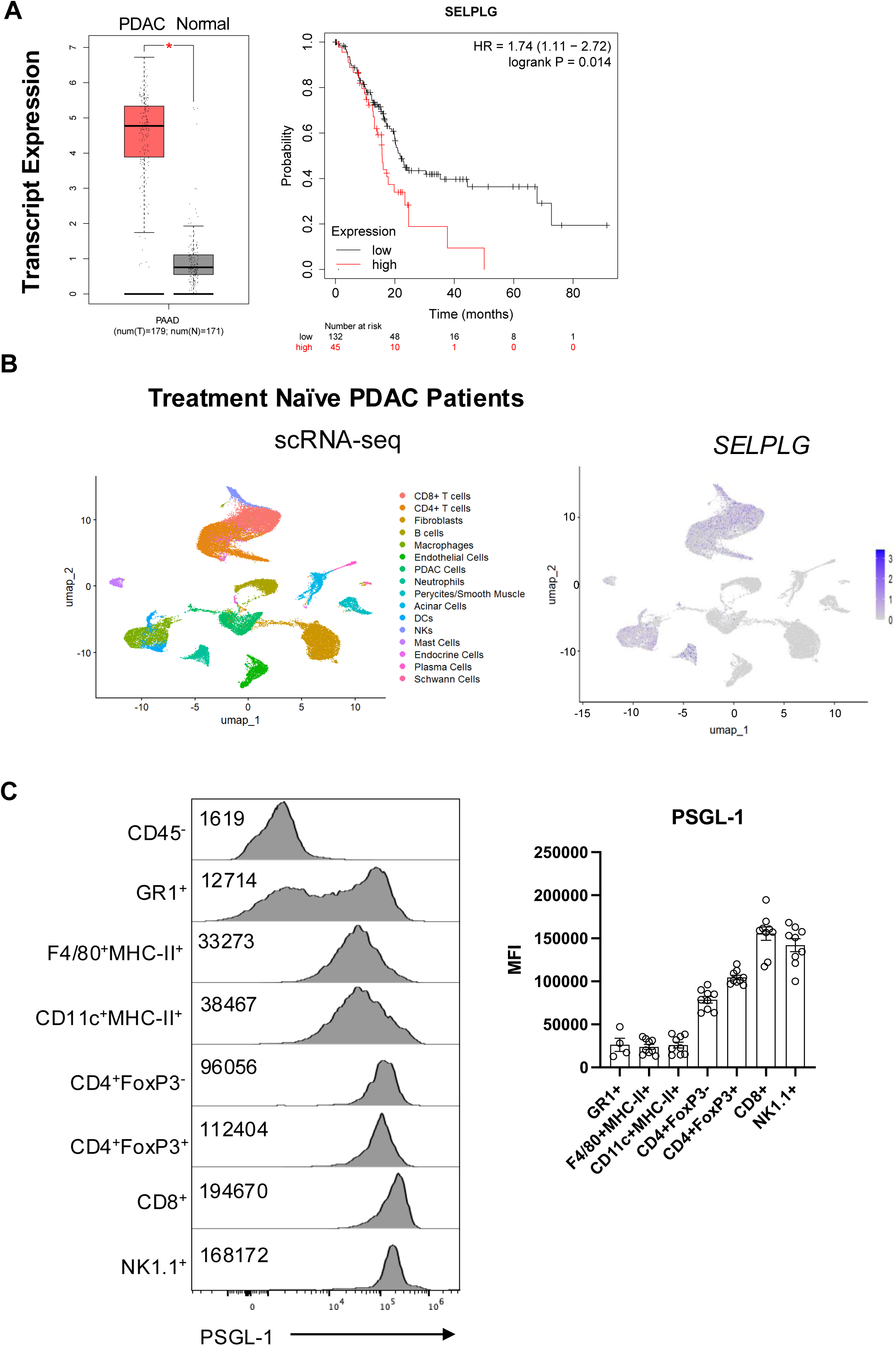
PSGL-1 is expressed by tumor-infiltrating immune cells in human and mouse PDAC tumors. **(A)** Box plot generated using GEPIA server and TCGA database evaluating SELPLG expression in pancreatic adenocarcinoma tumors (indicated here as PAAD) shown in red (n=179) paired to normal tissue (gray, n=171). PSGL-1 is overexpressed in PDAC patients, p<0.05 calculated using one-way ANOVA. Each dot represents a sample, transcripts per million (left). PDAC patient survival with SELPLG expression was demonstrated using K-M plotter (right). (B) Publicly available single-cell RNA-sequencing dataset (PMID: 36944944) representing six treatment-naïve PDAC patient tumor resections was reanalyzed to generate a UMAP (Seurat V4, 2000 most variable genes, 20 dimensions), identifying 16 Luvain Clusters which were annotated based on published markers (left). PSGL-1 gene (SELPLG) expression overlay on UMAP clusters (right). **(C)** Flow cytometry assessment of PSGL-1 expression on tumor-infiltrating immune cells in mouse orthotopic pancreatic cancer model KPC.4662. Representative histograms (left) and bar graph (right) of PSGL-1 MFI. Tumor immune infiltrates were evaluated on D26 after injection of 2.5×10^4^ KPC.4662 tumor cells into the pancreas tail. Data from 2 independent experiments, n=4-9 mice/group.

Our syngeneic orthotopic model uses KPC-derived PDAC cells harboring Kras^G12D^ and Trp53^R172H^ mutations—the latter being the murine equivalent of the common TP53^R175H^ mutation found in human PDAC. These mutations reflect the genetic landscape of PDAC, with *KRAS* mutations present in ∼85% and *TP53* mutations in ∼50% of patients. (Lee et al., 2016). This model recapitulates many features of human PDAC including limited representation of cytotoxic CD8^+^ T cells, greater representation of suppressive immune cells, and resistance to immunotherapies. To assess PSGL-1 expression in KPC-derived tumors, we implanted KPC cells orthotopically into the pancreata of WT (C57BL/6) mice. At 26 days after injection, we found PSGL-1 expression exclusively on CD45^+^ cells by flow cytometry. Within the CD45^+^ population, we show PSGL-1 expression on CD4^+^ and CD8^+^ T cells, NK cells, and myeloid cells (macrophages, dendritic cells, granulocytes) (**Fig. 1C**). CD8^+^ T cells and NK cells displayed the highest expression on a per cell basis (MFI) (**Fig. 1D**). These data indicate that, as with human PDAC tumors, immune cells become established in the TME but fail to restrict tumor growth.

### PDAC tumor growth is restrained by PSGL-1-deficiency

As PSGL-1 drives exhaustion of CD8^+^ T cells, we hypothesized that its absence would enhance anti-tumor immune responses by improving T_EFF_ functions (Tinoco et al., 2016). We found that at 28 days post-orthotopic implantation, tumor weights were reduced by approximately 50% in PSGL-1^-/-^ mice (1.138 g WT vs 0.46 g PSGL-1^-/-^, p < 0.0001) with 70% of PSGL-1^-/-^ mice demonstrating at least a 90% reduction in tumor size relative to WT mice (**Fig. 2A**). PSGL-1^-/-^ mice also demonstrated significantly decreased metastatic burden, as evaluated by detection of tumor clusters in the lung tissue (**Fig. 2B**). To validate this finding, we injected KPC cells intravenously and assessed colonization of the lungs. At 17 days post-injection, we found that KPC cell colonization in the lungs was significantly reduced in PSGL-1^-/-^ mice compared to WT controls (**Fig. 2C**), indicating a direct effect on metastatic outgrowth rather than an indirect consequence of primary tumor size. Our results support the notion that tumor reactive T cells can relocate to distal sites where they mediate anti-tumor responses.

**Figure 2.**
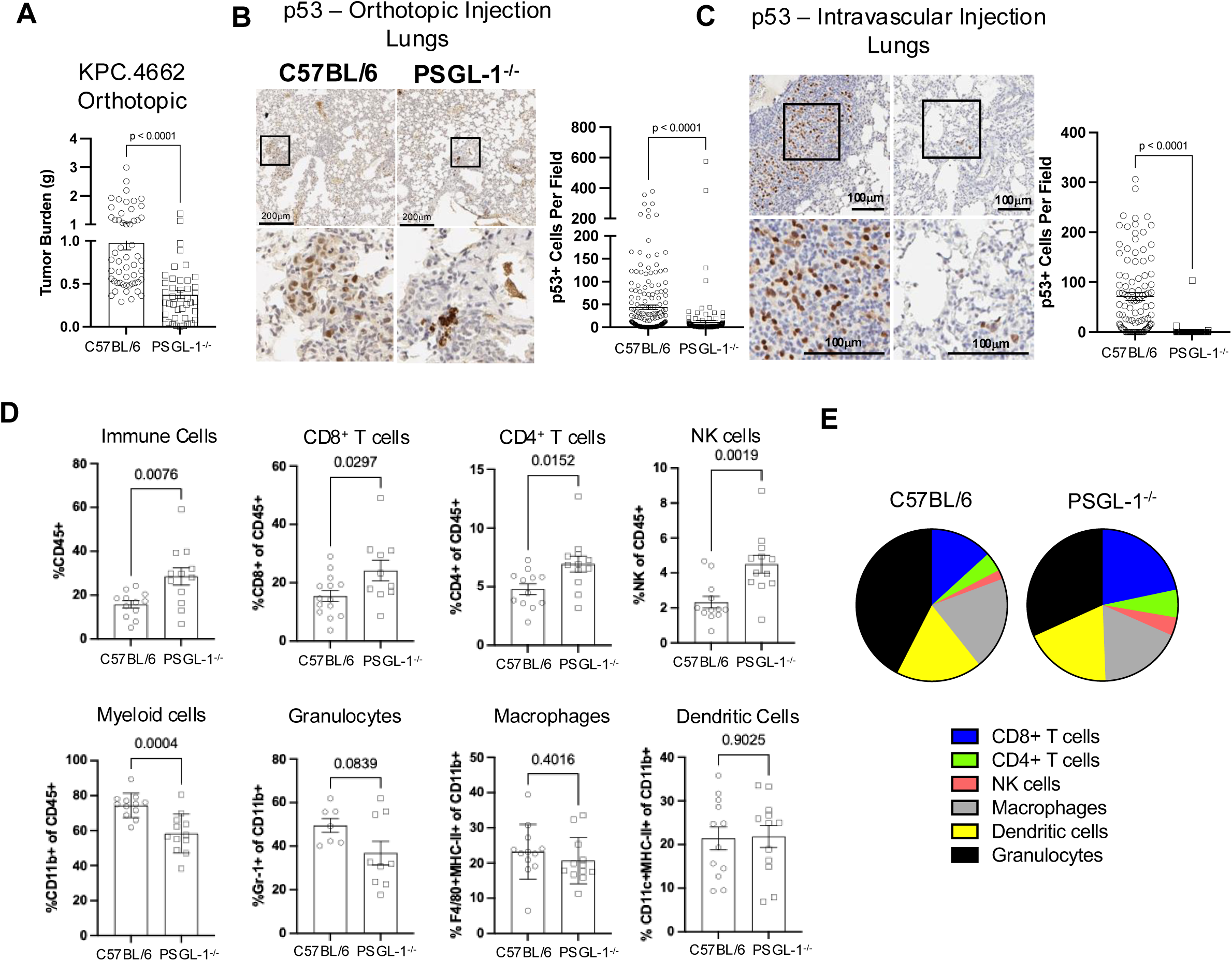
Immune deficiency of PSGL-1 promotes control of local and metastatic PDAC disease and alters intratumoral immune compartment distribution. For the orthotopic model, 2.5×10^4^ KPC.4662 tumor cells were injected into the pancreas tail of C57BL/6 control or PSGL-1^-/-^ mice and tumors were assessed on days 21-28 post injection. For metastasis model, 2×10^5^ KPC.4662 tumor cells were injected into the tail vein of C57BL/6 control or PSGL-1^-/-^ mice and tumors were assessed on day 17 post injection. **(A)** Dot plot/bar graph of tumor burden (primary + non-primary tumor weight, g) from orthotopic model on days 26-28 post injection. Data from >5 independent experiments, n=3-12 mice/group. **(B)** Representative images (left) and quantification (right) of p53^+^ cells from the lungs of mice 28 days post-orthotopic injection. **(C)** Representative images (left) and quantification (right) of p53^+^ cells from the lungs of mice 17 days post I.V. injection. Data in B and C representative of 2 independent experiments, n=10 mice per group, 10 fields captured per sample. **(D)** Frequencies of immune cell populations within orthotopic tumors 26 days post-injection as determined by flow cytometry. Each dot represents an individual mouse from 3 independent experiments. Data was normally distributed and assessed using an unpaired t test**. (E)** Pie chart depicting averaged flow cytometry data of immune cell populations as a percent of total CD45.2^+^ population in C57BL/6 versus PSGL-1^-/-^ mice.

The PDAC TME is complex and heterogeneous with a characteristic stromal barrier and desmoplasia formed in part by collagen-deposition by cancer-associated fibroblasts (CAFs) (Masugi, 2022). To further evaluate the effects of PSGL-1 deficiency, we assessed desmoplasia by measuring collagen deposition using trichrome staining of PDAC tumor sections. Tumors from PSGL-1^-/-^ mice showed only a minimal increase in collagen, suggesting that this aspect of CAF function remains largely intact in the absence of PSGL-1 (**Fig. S1A**).

We then investigated the effect of PSGL-1 deficiency on the representation of immune cell subsets (CD45^+^). Using flow cytometry to identify infiltrating immune cells, we observed an increased frequency of CD45^+^ immune cells in PDAC tumors from PSGL-1^-/-^ mice (**Fig. S1B, C**). In this population, we identified T cells (CD4^+^, CD8^+^), NK cells (NK1.1^+^), and myeloid cells (CD11b^+^). Of myeloid cells, we further distinguished granulocytes (Gr1^+^), macrophages (F4/80^+^, MHC-II^+^), and dendritic cells (CD11c^+^, MHC-II^+^). Decreased tumor size in PSGL-1^-/-^ mice was correlated with a greater abundance of CD8^+^ T cells, CD4^+^ T cells and NK cells (**Fig. 2D**), and of these lymphocyte subsets, CD8^+^ T cells had the highest representation (**Fig. 2E**). Conversely, there were fewer CD11b^+^ myeloid cells in PDAC tumors from PSGL-1^-/-^ mice, although we did not observe a significant decrease in any one myeloid compartment assayed (**Fig. 2D, E**). These data indicate that PSGL-1 is a key regulator of immune responses in PDAC, driving a shift in the tumor microenvironment from a non-inflamed, T cell–excluded state to one enriched with T cell infiltration.

### PSGL-1-deficiency promotes remodeling of the PDAC TME

Since we observed that PSGL-1-deficiency alters the representation of immune cells in the PDAC TME, we assessed their spatial organization within the tumors using multiplex immunofluorescence histology (CODEX, Akoya PhenoCycler). Using a validated antibody staining panel for mouse tissues, we compared the localization and distribution of immune cells within orthotopic tumors from WT and PSGL-1^-/-^ mice at 21 days post-injection, a timepoint at which the tumors from both cohorts were comparable in weight. We first evaluated the composition of the PDAC tumors with respect to the vasculature, tumor cells, immune cells, and CAFs (PDGFRα^+^) in all tumors by UMAP clustering (**Fig. 3A**, top panel). We then subdivided the immune cell subsets and proliferating vs. non-proliferating tumor cells by clustering (**Fig. 3A**, bottom panel), and assessed the relative proportions of these subsets within tumors from WT and PSGL-1^-/-^ mice, revealing notable shifts in their relative frequencies (**Fig. 3B**). For tumors from PSGL-1^-/-^ mice, most striking were a decrease in proliferating tumor cells (Ki67^+^) and an increase in CD8^+^ T cells and antigen-presenting cells (APCs)(see also **Fig. S2**). Increased proinflammatory myeloid subsets (Ly6C^+^ monocytes and CD68^+^ macrophages) were also observed when comparing tumors from WT and PSGL-1^-/-^ mice. In addition, the frequency of CAFs was diminished in these tumors. Heat map representation of the data (**Fig. 3C)** further underscores the differences in the composition of tumors from individual PSGL-1^-/-^ and WT mice and demonstrate the dramatic transformation of the spatial distribution of the cells (**Fig. 3D**).

**Figure 3.**
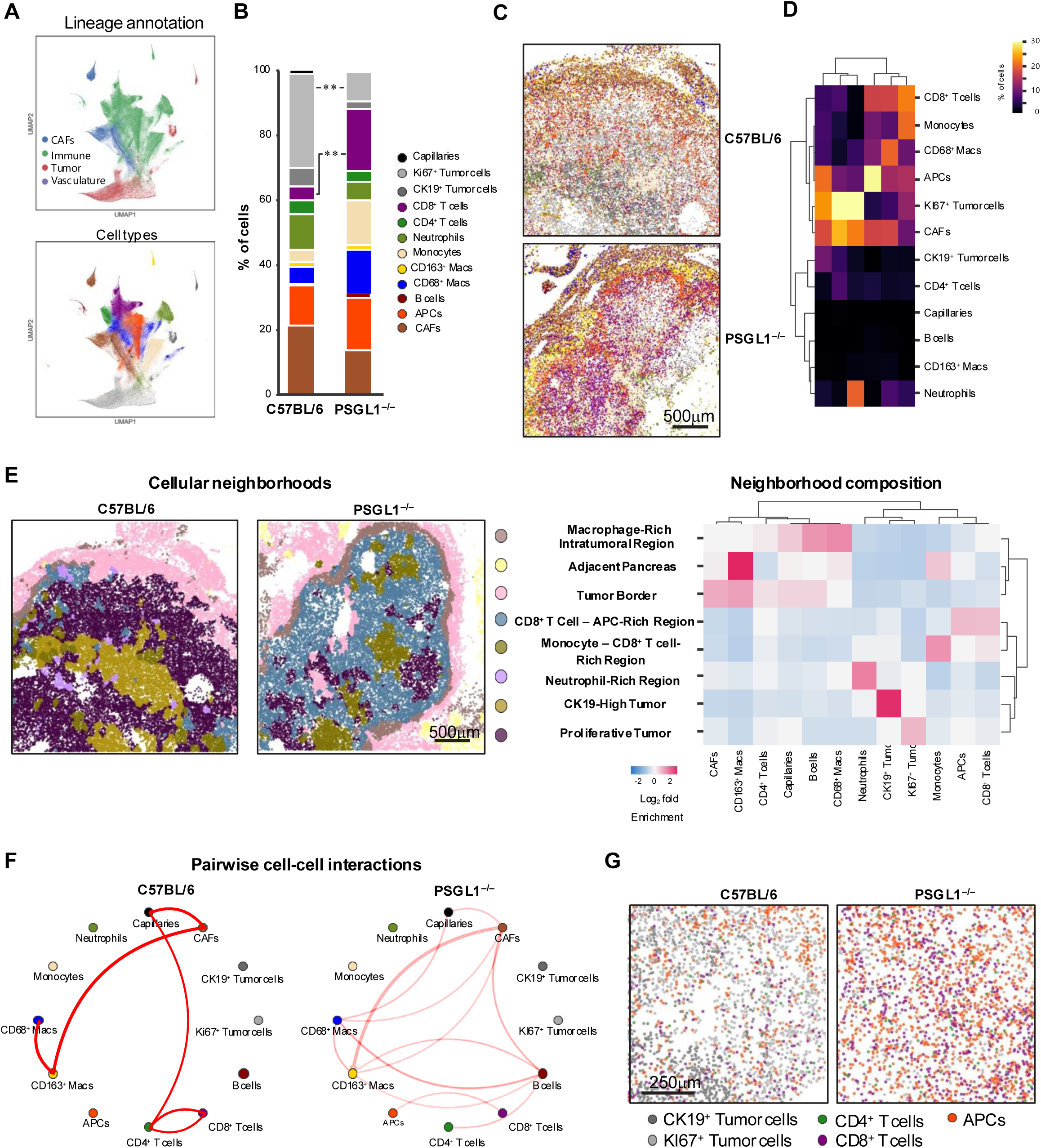
PSGL-1-deficiency promotes spatial remodeling of the PDAC tumor-immune microenvironment. Analysis of multiplexed immunofluorescence staining (Akoya PhenoCycler) on orthotopic tumors 21 days post-injection. Data from 3 independent experiments, n=3 mice/group. **(A)** Leiden clustering analysis of 451,636 single cells from tumors from C57BL/6 and PSGL-1^-/-^ mice. Clusters were manually annotated based on differential expression of cell type markers. **(B)** Relative distribution of cell types represented in KPC.4662 tumors from C57BL/6 or PSGL-1^-/-^ mice. **(C)** Representative images of KPC.4662 tumors from C57BL/6 or PSGL-1^-/-^ mice pseudocolored based on cell types as in A and B. D: Heat map of relative representation of the indicated cell types across 3 independent tumors per group. **(E)** Representative image of cellular neighborhoods (left) and enrichment of cellular populations within those neighborhoods (right) in tumors from C57BL/6 or PSGL-1^-/-^ mice. **(F)** Predicted pair-wise interaction networks of cells as evaluated in local cell neighborhood analyses (k=10). **(G)** Representative images showing co-localization of CK19^+^ tumor cells (non-proliferative) and Ki67^+^ tumor cells (proliferative), CD4^+^ and CD8^+^ T cells, and APCs in tumors from C57BL/6 and PSGL-1^-/-^ mice.

In addition to identifying cells that are represented within a tissue, multiplexed immunofluorescence staining provides insight into potential cellular interactions. Cellular neighborhood analysis defines regions of similar composition within the TME and uses the enrichment of individual cellular identities to define neighborhoods (**Fig. 3E**, left panel). Our analysis of PDAC tumors from WT and PSGL-1^-/-^ tumors identified 8 neighborhoods which were subsequently named based on the enriched populations (**Fig. 3E**, right panels). We observed the greatest increased representation of regions associated with CD8^+^ T cells and APCs that were present in tumors from PSGL-1^-/-^ mice but largely absent in WT mice. Localization of monocytes within these regions was also observed in the tumors from PSGL-1^-/-^ mice but not WT mice. This analysis further demonstrates dramatic differences in regions of proliferative tumor cells.

We then used these enriched neighborhoods within the TME to predict pair-wise interaction networks and changes resulting from PSGL-1-deficiency. This analysis revealed that the potential interactions of CD8^+^ T cells within tumors from PSGL-1^-/-^ mice were primarily with APCs (CD11c^+^MHC-II^+^)(**Fig. 3F**). Several additional interactions indicative of greater potential for cellular cross talk were also revealed. In tumors from both WT and PSGL-1^-/-^ mice, macrophages bearing the suppressive phenotypic marker, CD163, were predicted to associate with CAFs, and not with T cells. To gain further insight into the changes in the distribution of T cells, we compared their localization with tumor cells and APCs (**Fig. 3G**). These images underscore the remarkable remodeling of the PDAC TME in the absence of PSGL-1 that enabled formation of the CD8^+^ T cell−APC niche that is largely devoid of tumor cells, an outcome that was not observed in WT mice. These results support the concept that in these regions, CD8^+^ T cell responses elicited by local antigen presentation via APCs supports cytotoxic destruction of proliferating tumor cells.

### Control of PDAC tumors by PSGL-1^-/-^ mice is T cell dependent

Previous studies have found that a PDAC TME with a high number of effector CD4^+^ and CD8^+^ T cells and limited suppressor cells correlates with better patient survival (Carstens et al., 2017). Given the increased infiltration of T cells into PDAC tumors in PSGL-1^-/-^ animals, we asked whether T cells were the primary driving factor of tumor growth control. We performed depletion of CD4^+^ and CD8^+^ T cells prior to tumor implantation. Anti-CD4 and -CD8 delivered on days -2 and -1 prior to tumor implantation was sufficient to deplete all peripheral T cells from the mice (**Fig. S3A**). Whereas T cell depletion had no impact on orthotopic tumor growth in WT mice, it ablated the protective effect of PSGL-1-deficiency (**Fig. 4A**). These results demonstrate that T cells are the critical mediators of PDAC tumor growth control in PSGL-1^-/-^ mice.

**Figure 4.**
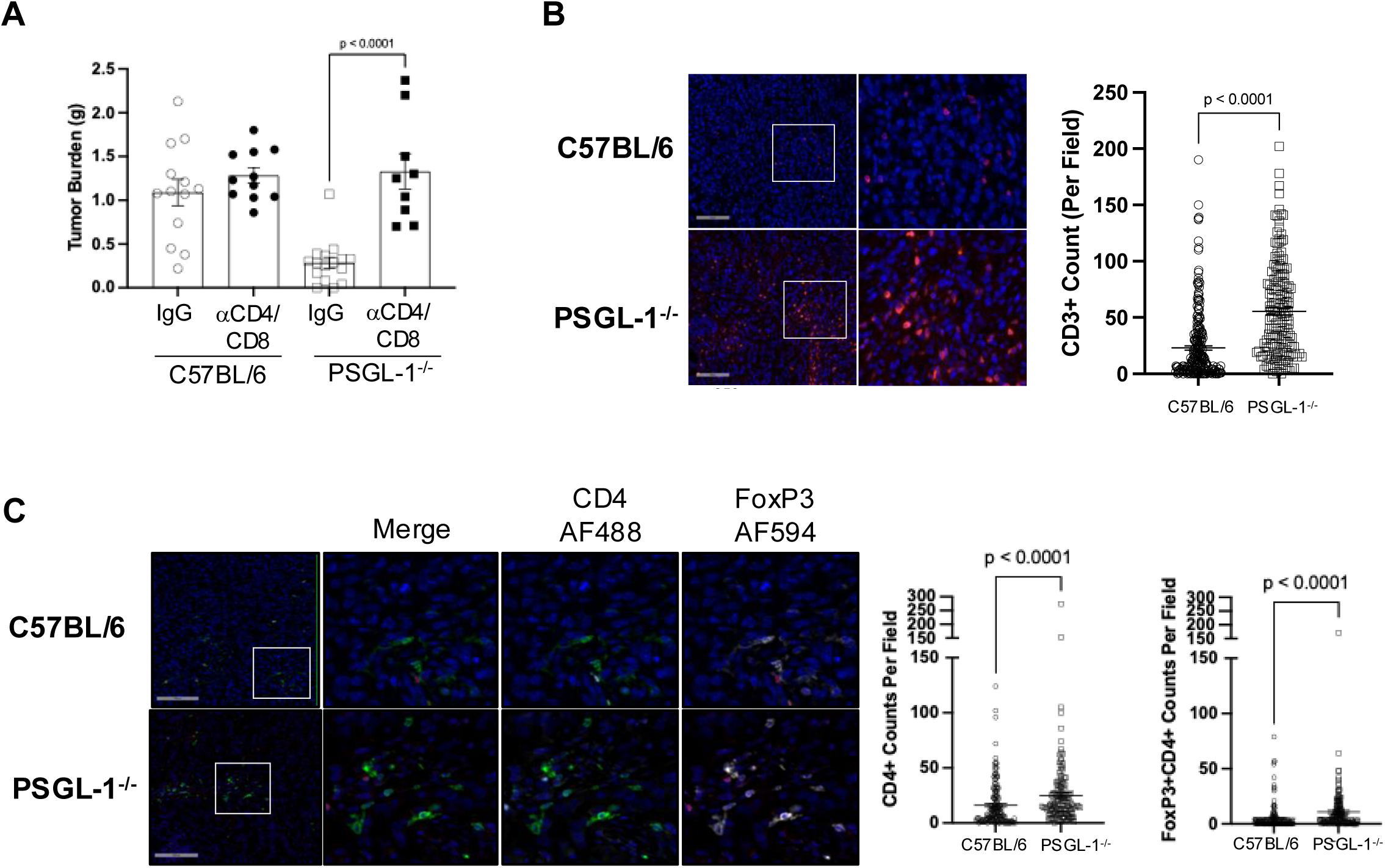
Control of PDAC tumor growth in PSGL-1^-/-^ mice is T cell-dependent. **(A)** CD4^+^ and CD8^+^ T cells were depleted from C57BL/6 and PSGL-1^-/-^ mice prior to the injection of 2.5×10^4^ KPC.4662 tumor cells into the pancreas tail. Tumors were assessed on days 26 post injection and compared to IgG-treated C57BL/6 and PSGL-1^-/-^ mice. Dot plot/bar graph of tumor burden (primary and non-primary tumor weight, g) from mice 26 days post injection. Data from 2 independent experiments, n=9-16 mice/group. Data was normally distributed except PSGL-1^-/-^ + IgG treatment. Statistics were assessed with an unpaired t test or Mann Whitney Exact test. (**B and C**) Orthotopic KPC.4662 tumors from C57BL/6 or PSGL-1^-/-^ mice were resected on day 28 post-injection, formalin fixed and paraffin embedded for sectioning and evaluation by immunofluorescent microscopy. Representative images of KPC.4662 tumor cores (exclusive of the outer 200μm of tumor) with DAPI and anti-CD3-AF564 staining **(B)** or CD4-AF488 and FoxP3-AF594 **(C)**. Data representative of 2 independent experiments. Data are non-normally distributed and were assessed using Mann Whitney Exact tests. B and C histograms each dot in quantification of cells represents cells per 400μm x 400μm field, 7-31 fields per mouse based on tumor size.

We next sought to understand differences in the T cell responses between WT and PSGL-1^-/-^ mice. Despite comparable desmoplasia in tumors from WT and PSGL-1^-/-^ mice, which can be a barrier to T cell infiltration into PDAC tumor cores, we observed that CD3^+^ T cells were significantly increased within intratumoral regions in PSGL-1^-/-^ mice (**Fig. 4B**). In these tumors, the numbers of CD4^+^ T cells were modestly increased as were the numbers of FoxP3^+^ T_REGs_ (**Fig. 4C**). We conclude that CD8^+^ T cells were the most significantly increased population among CD3^+^ T cells. Therefore, the ratio of conventional CD4^+^ and CD8^+^ T cells to FoxP3^+^ T_REGs_ is much higher in PSGL-1^-/-^ mice (0.36) compared to WT mice (0.13), a further indication of a greater anti-tumor T cell response and a capacity of PSGL-1^-/-^ T cells to withstand the hostile PDAC TME.

### PSGL-1 functions to regulate T cell differentiation and anti-tumor T cell responses to PDAC

Repeated exposure to antigenic stimulation results in the progressive loss of CD8^+^ T cell function and terminal differentiation. As there are no currently defined epitopes that can be used to evaluate antigen-specific CD8^+^ T cells in response to KPC-derived tumors, we evaluated the phenotype of CD44^hi^ CD8^+^ tumor-infiltrating T cells (**Fig. 5A**). Analysis of expression and co-expression of IRs associated with functional T cell exhaustion (PD-1, LAG3, TIM-3, 2B4, and CD160) identified that ∼60% of WT CD8^+^ T cells in the tumors co-expressed 3 or more of these markers while only ∼30% of PSGL-1^-/-^ CD8^+^ T cells co-expressed 3 or more markers; almost 50% of PSGL-1^-/-^ CD8^+^ T cells co-expressed only 2 markers (**Fig. 5B**). In comparison to WT, more PSGL-1^-/-^ CD8^+^ T cells in PDAC tumors had upregulated CX3CR1, a marker of effector CD8^+^ T cells in anti-tumor immunity (Zwijnenburg et al., 2023), and displayed lower expression of PD-1 (**Fig. 5C, D**, top panels). We additionally evaluated expression of CD38 and CD101, which when co-expressed demarcate terminally differentiated CD8^+^ T cells (Philip et al., 2017), and found significantly fewer co-expressing CD8^+^ T cells from PSGL-1^-/-^ compared to WT and decreased expression of CD101 (**Fig. 5C, D**, bottom panels). We also evaluated co-expression of TIM-3 and CD101, a highly proliferative transitory population of CD8^+^ T cells (Hudson et al., 2019). This population was significantly increased in PSGL-1^-/-^ compared to WT (**Fig. 5E, F**). The data show that PSGL-1 deficiency supports greater infiltration of CD8^+^ T cells that are less phenotypically exhausted.

**Figure 5.**
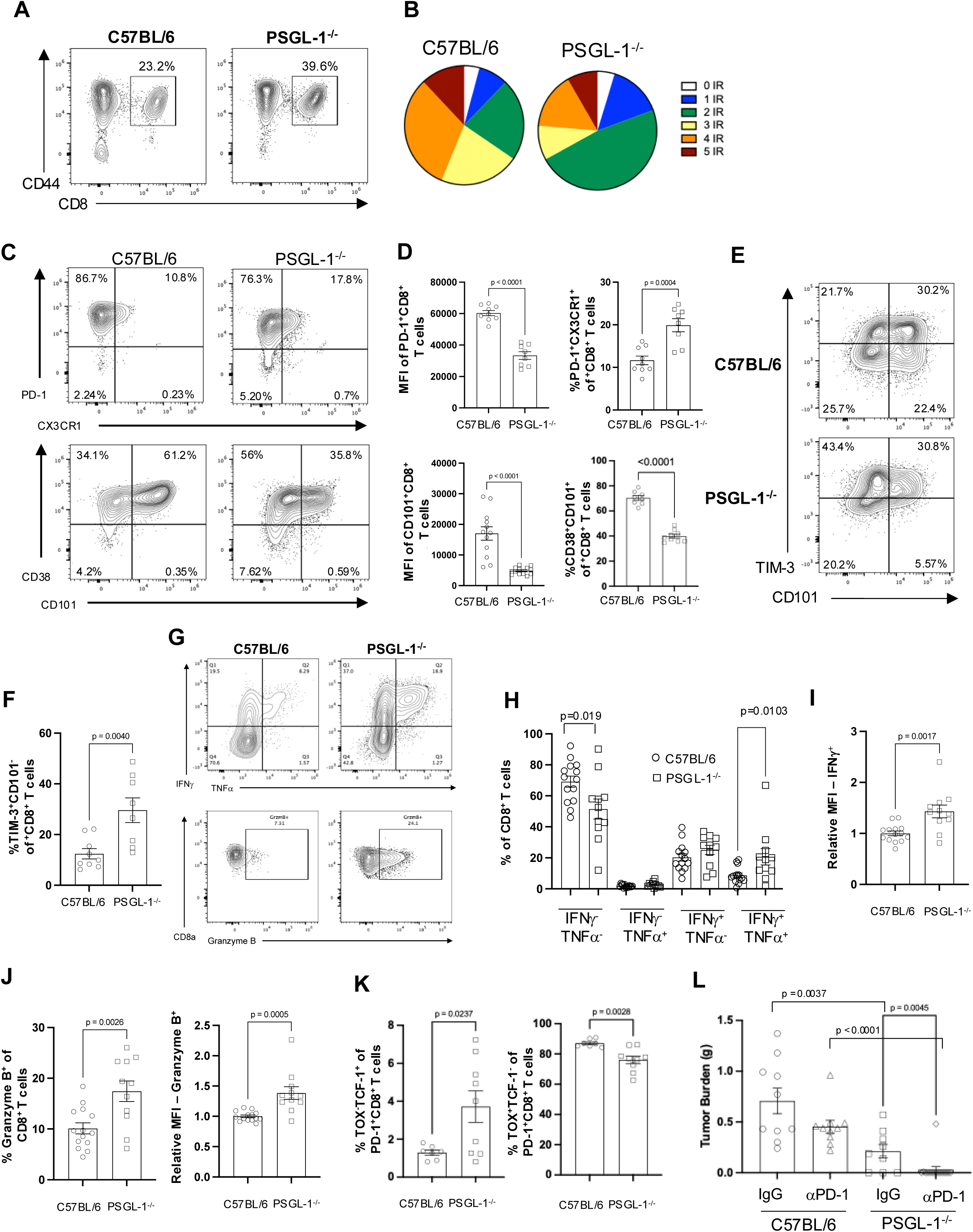
PSGL-1-deficiency synergizes with PD-1 immune checkpoint blockade to promote clearance of PDAC tumors. **(A)** Representative FACS plots of CD44 expression and CD8^+^ T cells within lymphocytes in orthotopic KPC.4662 tumors from C57BL/6 or PSGL-1^-/-^ mice on day 26 post-injection. **(B)** Pie graphs depicting co-expression of PD-1, LAG3, TIM-3, 2B4, and CD160 inhibitory receptors on intratumoral CD44^hi^ CD8^+^ T cells in C57BL/6 versus PSGL-1^-/-^ mice. **(C)** Representative FACS plots of PD-1 and CX3CR1 expression (top) and CD38/CD101 co-expression (bottom) on CD44^hi^ CD8^+^ T cells. **(D)** Quantification of median fluorescence intensity of PD-1 (top left) or CD101 (bottom left) in PD-1^+^ or CD101^+^ CD8^+^ T cells. Frequency of PD-1/CX3CR1 co-expressing (top right) or CD38/CD101 co-expression (bottom right) CD8^+^ T cells. **(E)** Representative FACS plots showing TIM-3 and CD101 expression on CD44^hi^ CD8^+^ T cells. **(F)** Frequency of TIM-3^+^CD101^-^ cells within CD44^hi^ CD8^+^ T cells. **(G)** Representative FACS plots showing cytokine production (top) and Granzyme B expression (bottom) by C57BL/6 or PSGL-1^-/-^ CD8^+^ T cells following restimulation with PMA/Ionomycin for 4 hours in the presence of Golgi inhibitors. **(H)** Quantification of cytokine-producing CD8+ T cells from G. **(I)** Relative MFI of IFNγ^+^ cells from G. **(J)** Quantification (left) and relative MFI (right) of Granzyme B-expressing CD8^+^ T cells from G. **(K)** Frequency of TCF-1 and TOX expressing intratumoral PD-1^+^CD44^hi^CD8+ T cells. A-K: Representative of at least 3 independent experiments; each dot represents an individual mouse. Data in D, F, and J are normally distributed and were assessed using an unpaired t test. Data in H are normally distributed except for double-positive cells; normally distributed data were assessed using an unpaired t test and non-normally distributed data was assessed using a Mann Whitney test. **(L)** KPC.4662-tumor bearing C57BL/6 and PSGL-1^-/-^ mice received 200μg of αPD-1 mAb (I.P.) or IgG antibody on days 10, 12, and 14 post-injection of tumors. Tumor burden was assessed on days 26. Data from 2 independent experiments, n=9-16 mice/group. IgG treatment group data was normally distributed; anti-PD-1 mAb treatment group was non-normally distributed. Statistics were assessed with an unpaired t test or Mann Whitney Exact test.

To address the functional potential of the intratumoral T cells, we evaluated cytokine production and Granzyme B (Gzmb) expression (**Fig. 5G-J**). Within PDAC tumors from PSGL-1^-/-^ mice, a much higher proportion of CD8^+^ T cells produced IFNγ, and coproduced IFNγ and TNFα when compared to CD8^+^ T cells within tumors from WT mice (**Fig. 5G, H**), and expressed increased IFNγ on a per-cell basis (**Fig. 5I**). In contrast, there were no differences in cytokine production by CD4^+^ T cells from the same mice (**Fig. S3B**). Consistent with improved cytotoxicity by CD8^+^ T cells, we found a greater frequency of PSGL-1^-/-^ CD8^+^ T cells expressing Gzmb than WT CD8^+^ T cells and more Gzmb per-cell (**Fig. 5G, J**). The presence of CD8^+^ T cells that expressed CX3CR1 in addition to polyfunctionality indicates an earlier state of T_EX_ differentiation wherein T_EFF_ functions are maintained (Raju et al., 2021).

To interrogate the transcriptional programs, we evaluated expression of the transcription factors TCF-1, which regulates stemness in CD8^+^ T cells (Zhang et al., 2021), and TOX, which determines the exhausted T cell fate of CD8^+^ T cells (Khan et al., 2019). Within intratumoral CD8^+^ T cells, we identified increased representation of TCF-1^+^ cells that did not co-express TOX in PDAC tumors from PSGL-1^-/-^ mice and, conversely, decreased representation of TOX^+^ cells that did not express TCF-1 CD8^+^ T cells (**Fig. 5K**). Thus, PSGL-1-deficiency restrains the development of T_EX_ within PDAC tumors to support their capacity for effector functions but also maintains TCF-1^+^ T_PEX_ CD8^+^ T cells. We conclude that greater CD8^+^ T cell infiltration of PDAC tumors in PSGL-1^-/-^ mice is accompanied by altered differentiation of T_EX_ subsets as shown by decreased expression of multiple IRs, decreased representation of cells expressing markers of terminal exhaustion, and maintenance of effector T cell functions together with a loss of proliferating tumor cells.

### PSGL-1-deficiency synergizes with PD-1 ICB to eradicate PDAC tumors

In patients, PD-1 ICB therapy designed to invigorate anti-tumor CD8^+^ T cell immunity fails to confer any benefit to PDAC patients. Similarly, KPC-derived tumors are resistant to anti-PD-1 treatment. Given that PSGL-1-deficiency promotes T cell infiltration into PDAC tumors and restrains differentiation of terminally exhausted CD8^+^ cells, we queried if PSGL-1^-/-^ CD8^+^ T cells would be responsive to therapy with PD-1 ICB. Orthotopic tumor-bearing WT and PSGL-1^-/-^ mice received three doses of anti-PD-1 ICB (days 10, 12, and 14) post-tumor cell injection. As expected, this treatment did not reduce primary tumor burden in WT mice, but remarkably, 85% of PSGL-1^-/-^ mice exhibited complete responses, with no detectable tumors 28 days post-implantation (**Fig. 5L**). The synergy between PSGL-1 deficiency and PD-1 blockade, leading to near-complete tumor eradication in PSGL-1^-/-^ mice, highlights the potential of targeting PSGL-1 to enhance anti-tumor immunity by promoting T_PEX_-to-T_EFF_ differentiation and overcoming resistance to PD-1 therapy— a major limitation in current PDAC treatment. These data suggest that PSGL-1 inhibition might also have potential for ameliorating PD-1 ICB resistance in other solid tumors.

Taken together, our data identify PSGL-1 as a pivotal regulator of CD8^+^ T cell exhaustion and immunosuppression in PDAC. Since PSGL-1 acts upstream of PD-1 and other IRs on T cells, it is notable that disabling this single IR in a PDAC model that is completely resistant to blockade of PD-1 and CTLA-4, afforded significant T cell-dependent tumor growth control that was accompanied by conversion to an inflamed TME as shown by significantly increased CD8^+^ T cell infiltration. The alteration in cellular composition and crosstalk with representation of CD8^+^ T cells adjacent to APCs in addition to much greater T_EFF_ functionality underscores the remarkable transformation of the anti-tumor CD8^+^ T cell response in the context of PSGL-1 deficiency. Such landmark changes in spatial distribution, differentiation, and function of CD8^+^ T cells have not been previously achieved in PDAC, and particularly by targeting a single inhibitory receptor.

Our finding of a greater abundance of TCF-1^+^ T_PEX_ CD8^+^ T cells in the TME of tumors from PSGL-1^-/-^ mice indicates that these cells can migrate into PDAC tumors as was previously described for other solid tumors (Fang et al., 2022; Siddiqui et al., 2019). Together with phenotypic and functional analyses, our data support the hypothesis that PSGL-1-deficiency restrains terminal differentiation of CD8^+^ T cells in response to PDAC to support the maintenance of these critical CD8^+^ T cell subsets within the TME. Future studies will be needed to determine the factors that drive altered cellular composition and spatial reorganization of the TME in the context of PSGL-1 deficiency. While our previous studies identified a direct role for PSGL-1 in regulating T_EX_ differentiation, the current findings reveal that PSGL-1 also modulates non–T cell immune responses within the PDAC TME—directly or indirectly—highlighting the need to further explore how CD8^+^ T cells coordinate with other immune populations. Together, these results demonstrate that loss of PSGL-1 signaling reprograms the TME toward a proinflammatory state that augments CD8^+^ T cell–mediated anti-tumor immunity. We conclude that inhibition of PSGL-1 signaling transforms the non-responsive TME of PDAC into one that enables mobilization of less exhausted CD8^+^ T cells with capacity for T_EFF_ functions and sensitivity to immunotherapy. Our work underscores the importance of focused development of PSGL-1-specific inhibitors and their evaluation in combination with existing immunotherapies.

## Acknowledgements

This work was supported in part by the Sanford Burnham Prebys NCI-Designated Cancer Center Support Grant, P30 CA030199. We would like to acknowledge the contributions of the following Sanford Burnham Prebys Core facilities: the Flow Cytometry Core (RRID:SCR_014854; Yoav Altman, Benji Portillo), the Vivarium Core (Buddy Charbono and Andy Vasquez), and the Histology Core (Guillermina Garcia and Monica Sevilla). This work was supported by The NIH Shared Instrumentation Grant S10OD032325 for the Cytek Aurora housed in the Flow Cytometry Core at Sanford Burnham Prebys.

## Author Contributions

J. Hope, Y. Zhang, H. Hetrick, E. S. Sanchez Hernandez, and S. Roy completed orthotopic tumor injections. J. Hope, H. Hetrick, E. S. Sanchez-Hernandez, M. Lin, S. Roy, A. Palete, and D. Otero completed sample processing and flow cytometry. B. Silvestri and Y. X. Wang completed PhenoCycler staining and analysis. A. Palete, H. Hetrick, and M. Lin completed mouse breeding and genotyping. S. Maganti and L. Ling completed histology staining. B. Smith and S. H. Nakil completed histology quantification. G. Romano completed scRNA-sequencing analysis. K. Byrne and E.S. Sanchez-Hernandez completed bulk RNA-sequencing analysis. J. Hope, Y. Zhang, C. Commisso, and L. Bradley designed the experiments and completed data analysis. J. Hope, H. Hetrick, and L. Bradley wrote the manuscript. All authors reviewed, critiqued, and approved the manuscript prior to submission.

## Data Availability Statement

The datasets generated or analyzed during the current study are available from the corresponding author on reasonable request.

## Methods

### Analysis of RNA data from human PDAC tumors

For profiling *SELPLG* mRNA expression in PDAC tumor and normal tissue, we used GEPIA (http://gepia.cancer-pku.cn/index.html), an online server with big data cancer analytics platform that allows the analysis of RNA sequencing expression data of tumors and normal tissue samples based on TCGA and the genotype tissue expression (GTEx). For Kaplan-Meier Analysis we used K-M plotter (https://www.kmplot.com/analysis/) (Nagy et al., 2021), a web tool was used to assess the association between *SELPLG* mRNA levels and overall survival (OS) of PDAC patients.

Single-cell RNA sequencing data were retrieved from GEO database entry GSE212966 (12 samples, 6 PDAC tumors, 6 tumor-adjacent samples). The data from all 12 samples were combined in R (version 4.1.2) using the Read10X() function from the Seurat package (4.1.4), and an aggregate Seurat object was generated (Fang et al., 2022; Hao et al., 2021; Hu et al., 2022; Stuart et al., 2019; Wang et al., 2021). Filtering was conducted by retaining cells with unique molecular identifiers (UMIs) greater than 200, expressed between 400 and 5000 genes, and mitochondrial content less than 10 percent. No sample batch correction was performed. This resulted in a total of 39,129 cells. Data were normalized using the “LogNormalize” method and a scale factor of 10,000. To reduce the dimensionality of this dataset, the 2000 most variably expressed genes were summarized by principle component analysis (PCA), and the first 20 principle components were further summarized using UMAP dimensionality reduction using the RunUMAP() function. Clustering was conducted with the FindClusters() function using 20 PCA components and a resolution parameter set to 0.8. The original Louvain algorithm was utilized for modularity optimization. The resulting 28 Louvain clusters were visualized in a two-dimensional UMAP representation and were annotated to known biological cell types using canonical marker genes (Supplementary Table 1). The top 10 genes per cluster were used to assign them to a biological cell type. These genes were annotated with data generated from the FindAllMarkers() Seurat differential expression analysis using the default two-sided non-parametric Wilcoxon rank sum test with Bonferroni correction using all genes in the dataset. To generate the violin plots VlnPlot() function was used. Additional packages used for data analysis include: ggplot2 (3.4.3), dplyr (1.1.2), SingleR (1.8.1), SingleCellExperiment (1.16.0), tidyverse (2.0.0).

### Mice

C57BL/6J and B6.Cg-PSGL-1*^tm1Fur^*/J (PSGL-1*^-/-^*) were obtained from the Jackson Laboratory. PSGL-1*^-/-^* were backcrossed to the Thy1.1 background (B6.PL-*Thy1^a^/*CyJ). All mice were kept in a barrier facility (certified by the Association for the Assessment and Accreditation of Laboratory Animal Care) on acidic water at the Sanford Burnham Prebys Medical Discovery Research Institute. All lines in the Bradley mouse colony are backcrossed to parental C57BL/6J or B6.PL-*Thy1^a^/*CyJ lines biannually. This study was carried out in accordance with the recommendations and approval of the Institutional Animal Care and Use Committee (IACUC).

### Tumor Cell Culture and Orthotopic Injection

LSL-Kras^G12-D/+^; Trp53^fl/+^; Pdx1-Cre (KPC) mice-derived PDAC cells (KPC.4662) were from Dr. Robert Vonderheide (University of Pennsylvania). KPC.4662 cells were cultured in DMEM supplemented with 10% fetal bovine serum (FBS, Corning), 10^5^ I.U. Penicillin, 10^5^ µg/mL Streptomycin, and 292 mg L-glutamine (Corning Cellgro). Cells were maintained *in vitro* at 37°C, 5% CO_2_. For *in vivo* experiments, tumor cell aliquots from the same passage were thawed and passaged 3 times prior to injection. For orthotopic tumor initiation, 6–8-week-old male or female mice were anesthetized with 100 mg/kg Ketamine and 20 mg/kg Xylazine and 2.5×10^4^ KPC.4662 cells suspended in PBS with 50% Matrigel (Corning) were injected into the tail of the pancreas. Tumors were harvested 3-4 weeks later for *ex vivo* analysis. For evaluation of metastasis development, 2×10^5^ KPC.4662 tumor cells suspended in 1X PBS were injected intravascularly into the tail vein of mice and assessed on day 17 post-injection.

### *In vivo* antibody treatments

For T cell depletion studies, C57BL/6 and PSGL-1^-/-^ mice received an I.P. injection of 0.5 mg of Rat IgG Isotype (Jackson ImmunoResearch), anti-CD8a (53-6.7, BioXCell), or anti-CD4 (GK1.5, BioXCell) monoclonal antibodies on days -2 and -1 prior to tumor cell injection. For PD-1 ICB, C57BL/6 and PSGL-1^-/-^ KPC.4662 tumor-bearing mice received an I.P. injection of 0.2 mg of Rat IgG Isotype (Jackson ImmunoResearch) or anti-PD-1 (RMP1-14, BioXCell) monoclonal antibodies on days 10, 12, and 14 post-tumor injection.

### Multiplexed immunofluorescence analysis of PDAC tumors

#### Tissue preparation and staining

PDAC tumors from WT and PSGL-1^-/-^ mice 21 days after orthotopic injection were dissected and embedded as described (Wang et al., 2022). Briefly, tumors were placed in OCT and quickly frozen by immersion in semi-frozen, liquid nitrogen-cooled isopentane. Tissue sections (10 um cryosection) were thawed in Hydration Buffer (S1) for 2 min and washed twice. Subsequently, sections were fixed in S1-diluted 1.6% PFA for 10 min and washed with S1 three times. 5 min washed was performed with S2 buffer and then the Antibody Mix 1 was added and incubated overnight at 4 °C in a hydration chamber. 0.2% gelatin-coated 22×22mm coverslips were washed twice with S2 buffer and fixed in S1-diluted 1.6% PFA for 10 min. Three washes with PBS were then performed and a final fixation step with 200 mg/ml BS3 (bis(sulfosuccinimidyl)suberate; Thermo Fisher) in anhydrous DMSO (Sigma) for 20 min. the coverslips were again washed three times with PBS and stored at 4 °C in Storage Solution (S4) until CODEX imaging. Antibody staining was performed using Antibody Mix 1 by adding: Ly-6G, CD11b, Ly-6C, F4/80, CD68, CD163, CD11c, MHCII, CD3, CD4, CD8a, CD27, TCRβ, CD90, PD-1, PDL-1, CD56, CD127, B220, CD19, Ki-67, PDGFRa, ERTR7, FN, Pan-lam, CD45.2, CD31, PSGL-1, FOXP3, CD103, aSMA, CK-19, CX3CR1, TIM-3, LAG-3 antibodies into in S2 buffer together with 5 uL of each of 1 mg/ml mouse IgG (Sigma), 1 mg/ml rat IgG (Sigma), Sheared salmon sperm DNA (10 mg/ml in H2O; Thermo Fisher), Mixture of non-modified CODEX oligonucleotides at a final concentration of 0.5 mM in TE buffer (IDT) solutions.CODEX was performed on a Phenocycler Open system (Akoya) on a Keyence BZ-810 automated microscope in 23 cycles. Fluorescent probes (5 uL of a 10 mM stock in 250 uL of plate buffer) for the matching antibody barcode was loaded into a 96-well plate as per Phenocycler protocol. The entire tumor section was captured in tiles with optical sectioning (3 x 2.4 um z-slices). For each CODEX experiment, a wild-type sample and a PSGL1^-/-^ sample were sectioned on a common coverslip, stained and imaged together.

#### Quantification and analysis

Imaging data was processed using CRISP-CODEX processor (Wang et al., 2022) to register (across image tiles and imaging cycles), deconvolve, and generate stitched extended-depth-of-field images of the entire tumor section. Cells were segmented using a modified version of CellSeg (Lee et al., 2022) that has been optimized for CRISP output (https://github.com/will-yx/CellSeg-CRISP). Quantified per cell staining intensity from images were pre-processed using HFcluster (Wang et al., 2022; https://github.com/will-yx/HFcluster/tree/anndata) for analysis in Scanpy (Wolf et al., 2018). Harmony was used to integrate replicate data across 3 experiments. Cell types were clustered by expression similarity using Leiden (resolution=2) and manually annotated by marker expression. Cell neighborhoods (Schürch et al., 2020) were determined by the composition of each cell and its nearest 40 neighbors, then clustered based on the relative abundance of each cell type (Leiden, resolution=0.2) which yielded 12 neighborhoods. Each neighborhood was then annotated based on enriched cell types and its spatial localization in the PDAC tumor. Cell type enrichment was calculated as log2fold enrichment of its abundance in each neighborhood over the total abundance in the whole tissue.

### Histology

Tumor sections were fixed with 10% buffered zinc formalin (StatLab Medical Products) for 24 hours, washed with 1X PBS, and stored in 70% EtOH until standard paraffin embedding and sectioning. Following deparaffinization and rehydration, heat-induced antigen retrieval on sections was performed using 0.01 mol/L citrate buffer (pH 6.0) and a microwave. Sections were then rinsed three times with 1X PBS and immersed in 1% hydrogen peroxide for 30 minutes at room temperature before permeabilization with 0.1% Triton-X100 in TBS containing 0.1% Tween 20 (TBST) for 10 minutes. Sections were blocked with 10% goat serum and 1% BSA in TBST for 1 hour. Sections were stained with the primary antibodies indicated below overnight at 4°C, followed by incubation at RT with the indicated secondary antibodies. Nuclear counterstaining was performed using hematoxylin and sections were then dehydrated and mounted with coverslips using Permount mounting medium (Thermo Fisher Scientific). Immunofluorescence and H&E staining was performed by the Sanford Burnham Prebys Medical Discovery Institute Histology Core facility. p53 staining was completed by the Commisso lab. Images were captured using brightfield microscope installed with an INFINITY camera (Lumenera). Quantification was performed using FIJI.

### Flow Cytometry

Tumors were processed using the Miltenyi Mouse Tumor Dissociation Kit and gentleMACS C tubes (Miltenyi) according to manufacturer’s instructions, except with the intentional exclusion of enzyme R (due to cleavage of PSGL-1 and CD44). Tumors were treated with RBC lysis buffer (0.145 M NH_4_Cl, 0.05 M Tris-HCl, pH 7.2) for ∼1 minute at room temperature. Processed tumors were maintained in RPMI 1640 (Corning) supplemented with 5% FBS, 1% Penicillin/Streptomycin/L-glutathione and 75 ug/mL DNAse I (Worthington). In all stains, cells were pre-treated with 2.5 µg/mL anti-CD16/32 (Fc Block; 2.4G2; BioLegend, San Diego, CA) for 15 minutes before continuing with surface staining. For surface stains, 2×10^6^ cells were stained for 20 min on ice. Cells were stained with the fluorochrome conjugated monoclonal antibodies as indicated below. After staining, cells were washed twice with 1X HBSS containing 3% FBS and 0.02% sodium azide and fixed with 1% formaldehyde. For staining of intracellular transcription factors, 2×10^6^ cells were stained as above for surface markers. Following the surface stain, cells were permeabilized with eBioscience FoxP3 fixation buffer (ThermoFisher) overnight at 4°C, washed using eBioscience 1X Perm Wash (ThermoFisher), followed by intracellular staining for 45 minutes. Cells were then washed 2 times with 1X Perm Wash containing 0.02% sodium azide and fixed with 1% formaldehyde. Cells were stained with the fluorochrome conjugated monoclonal antibodies are indicated below. For staining of intracellular cytokines, 2×10^6^ cells were stimulated with anti-CD3 (145-2C11, BioXCell) overnight at 37°C, 5% CO_2_ in the presence of GolgiPlug (BD Biosciences), Monensin (BioLegend), and 10 U/mL recombinant human IL-2. Cells were surface stained as above then fixed overnight at 4°C with FoxP3 fixation buffer (eBioscience), washed using Perm/Wash buffer (eBioscience) and stained for intracellular cytokines for 45 minutes at 4°C. Cells were stained with the fluorochrome conjugated monoclonal antibodies indicated in the table below. 1X Perm Wash containing 0.02% sodium azide and fixed with 1% formaldehyde. Data was acquired on a LSRFortessa X-20 (BD) or 5-laser Aurora (Cytek) and analyzed using FlowJo v10 software (BD).

#### Statistics

For *in vivo* studies, a power analysis was performed to determine the number of animals per group per experiment, assuming 80% power and two-sided 0.05 significance. For each experiment, the number of animals per group are indicated in the figure legend. The number of replicate experiments is noted in each figure legend. Specific statistical analyses for big data sets are indicated in their specific methods section. For flow cytometry data, the normality of the population distribution was assessed using the Shapiro-Wilk normality test by GraphPad Prism 10. Significant differences between normally distributed populations were assessed using a two-tailed, unpaired *t*-test; significant differences between non-normally distributed populations were assessed using a two-tailed Mann Whitney exact test. The tests performed are denoted in each figure legend and subsequent p-values are annotated in the associated figure. Error bars denote standard error of the mean.

### Antibodies

#### Histology

Primary antibodies: CD3(DF 1:100, RRID: AB_443425), CD4(DF 1:200, RRID: AB_2686917), Ki67 (DF 1:200, RRID: AB_302459), FoxP3(DF 1:50, RRID: AB_467576), P53(DF 1:500, RRID: AB_331743).

#### CODEX

LY6G(DF 1:100, RRID: AB_1089180), CD11b (DF 1:100, RRID: AB_312785), Ly6C (DF 1:100, RRID: AB_2563783), F4/80 (DF 1:100, RRID: AB_893506), CD68 (DF 1:100, RRID: AB_2044004), CD163 (DF 1:100, RRID: AB_2716934), CD11c (DF 1:100, RRID: AB_313771), MHCII(DF 1:100, RRID: AB_313317), CD3 (DF 1:50, RRID: AB_312659), CD4 (DF 1:50, RRID: AB_312687), CD8a (DF 1:50, RRID: AB_314120), CD27 (DF 1:25, RRID: AB_2561786), TCRB (DF 1:100, RRID: AB_313425), CD90 (DF 1:100, RRID: AB_313169), PD1 (DF 1:100, RRID: AB_2941806), PDL1 (DF 1:100, RRID: AB_2773715), CD127 (DF 1:25, RRID: AB_1937287), B220 (DF 1:50, RRID: AB_312987), CD19 (DF 1:100, RRID: AB_313637), Ki67 (DF 1:100, RRID:AB_10854564), PDGFRa (DF 1:100, RRID: AB_2236897), ERTR7 (DF 1:100, RRID: AB_915429) Fn1 (DF 1:100, RRID: AB_447655), Pan-Laminin (DF 1:100, RRID: AB_477163), CD45.2 (DF 1:100, RRID: AB_2563751), CD31 (DF 1:100, RRID: AB_312897), FoxP3 (DF 1:100, RRID: AB_439745), alpha SMA (DF 1:100, RRID: AB_2572996), PSGL1 (DF 1:100, RRID: AB_10950305), CD103 (DF 1:50, RRID: AB_535945), Cytokeratin, pan (DF 1:100, RRID: AB_3284598), CX3CR1 (DF 1:100, RRID: AB_2564313), TIM3 (DF 1:100, RRID : AB_10949464), LAG3 (DF 1:100, RRID: AB_10949602), rPSGL1 (DF 1:25, custom made, Aragen).

#### Flow Cytometry

Spark UV387 CD45.2 (DF 1:200, RRID: AB_2941386), BUV737 PSGL1 (DF 1:200, RRID: AB_2871141), BV421 CD11c (DF 1:200, RRID: AB_11219593), BV510 CD4 (DF 1:400, RRID: AB_2564587), BUV395 CD4 (DF 1:200, RRID: AB_2738426), BV605 NK-1.1 (DF 1:200, RRID: AB_2562274), BV650 GR-1 (DF 1:200, RRID: AB_2686974), BV711 I-A/I-E, (DF 1:200, RRID: AB_2565976), BV785 CD11b (DF 1:400, RRID: AB_2561373), Apotracker Green (DF 1:200, Biolegend 427403), PerCP/Cyanine5.5 CD8a (DF 1:500, RRID: AB_2075238), PE FoxP3 (DF 1:200, RRID: AB_1089117), PE/Cyanine7 F4/80 (DF 1:400, RRID: AB_893478), AlexaFluor700 CD44 (DF 1:400, RRID: AB_493713), BUV496 CD44 (DF 1:200, RRID: AB_2870671), PerCPCy5.5 CD90.1 (DF 1:500, RRID: AB_961437), PerCPCy5.5 CD90.2 (DF 1:500, RRID: AB_2571945), APC CD90.2 (DF 1:500, RRID: AB_313183), BUV395 CD8a (DF 1:200, RRID: AB_2732919), BV605 CD8 (DF 1:200, RRID: AB_2562609), BV785 PD-1 (DF 1:200, RRID: AB_2563680), BV421CX3CR1 (DF 1:200, RRID: AB_2565706), APC/Fire750 CD38 (DF 1:200, RRID: AB_2876402), PECy7 CD101 (DF 1:200, RRID: AB_2573378), BV605 TIM3 (DF 1:200, RRID: AB_2616907), BV711 TIM3 (DF 1:200, RRID: AB_2716208), APC IFNg (DF 1:200, RRID: AB_315404), PECy7 TNFa (DF 1:200, RRID: AB_2256076), PE Granzyme B (DF 1:200, RRID: AB_10561690), ef660 TOX (DF 1:100, RRID: AB_2574265), PE TCF1(DF 1:200, RRID: AB_2687845)

## Supplemental Figures for

**Supplemental Figure 1.**
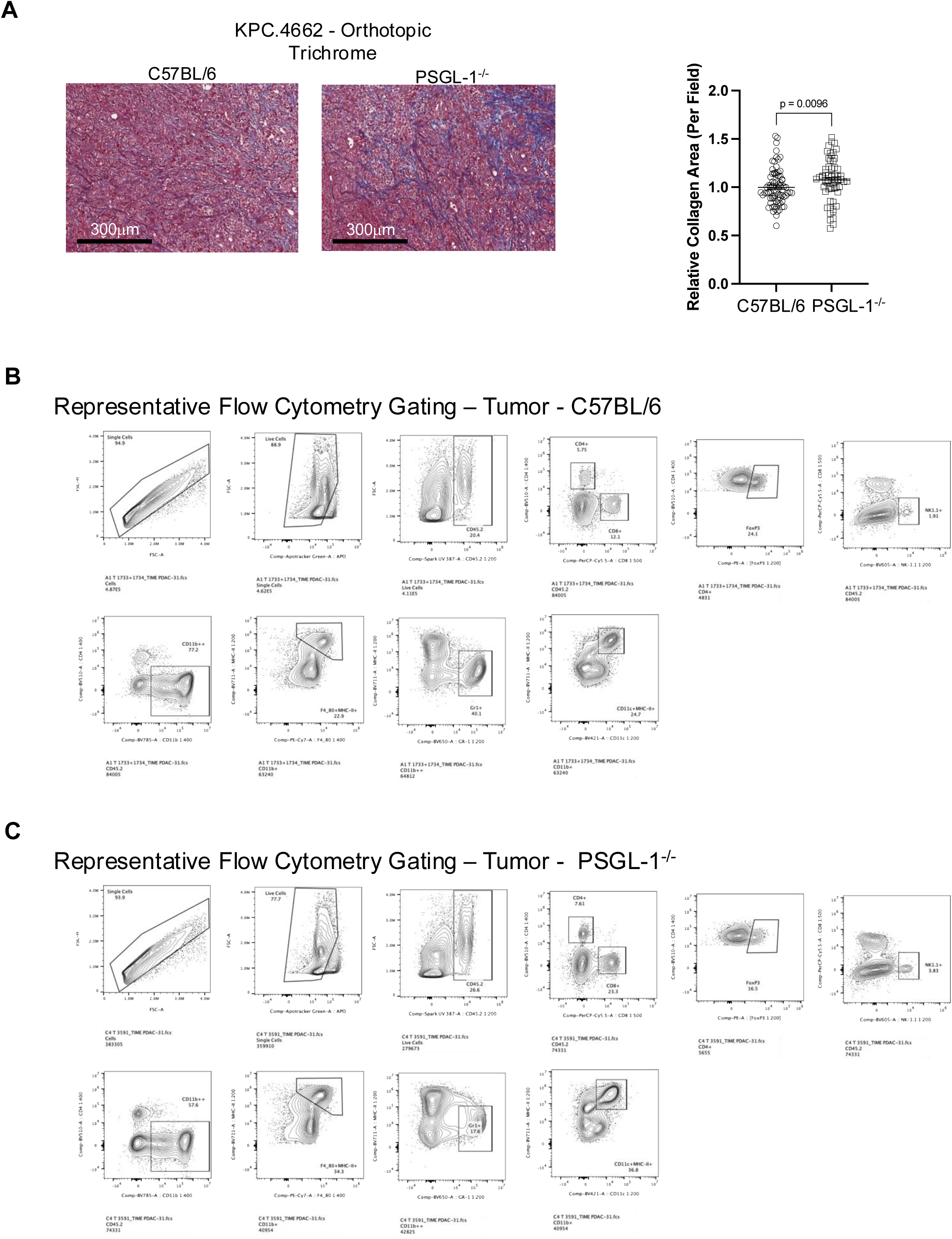
Analysis of KPC.4662 tumors and immune infiltrates in C57BL/6 and PSGL-1^-/-^ mice. **(A)** KPC.4662 tumors from C57BL/6 and PSGL-1^-/-^ mice on day 28 post-injection were assessed for collagen deposition by trichrome staining. Left: representative trichrome staining; right: relative collagen area quantification. **(B and C)** Representative flow cytometry gating strategy to identify immune populations within single-cell suspensions generated from C57BL/6 (B) or PSGL-1^-/-^ mice.

**Supplemental Figure 2.**
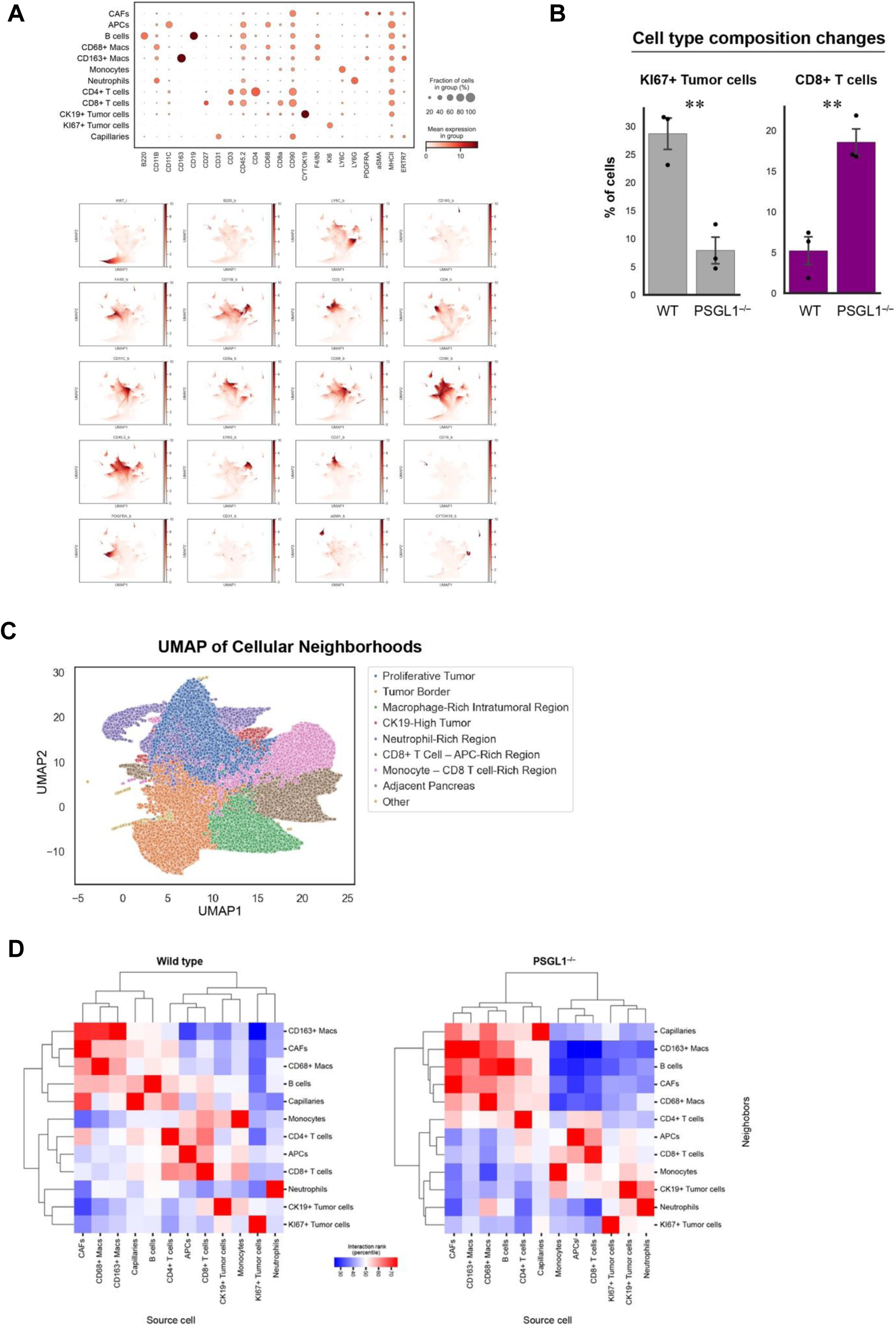
PhenoCycler analysis of KPC.4662 tumors in C57BL/6 and PSGL-1^-/-^ mice. **(A)** (Top) Bubble plot showing the mean expression of each marker and fraction of cells expressing each marker within the identified cellular populations. (Bottom) Density plot showing the localization of marker expression within the UMAP. **(B)** Dot plot/bar graph showing changes in proliferating tumor cells and CD8^+^ T cells within the individual tumors from C57BL/6 and PSGL-1^-/-^ mice. **(C)** UMAP of cellular neighborhoods defined within tumors from C57BL/6 and PSGL-1^-/-^ mice. **(D)** Heat maps of the predicted cellular interactions as determined by next-nearest neighbor analyses.

**Supplemental Figure 3.**
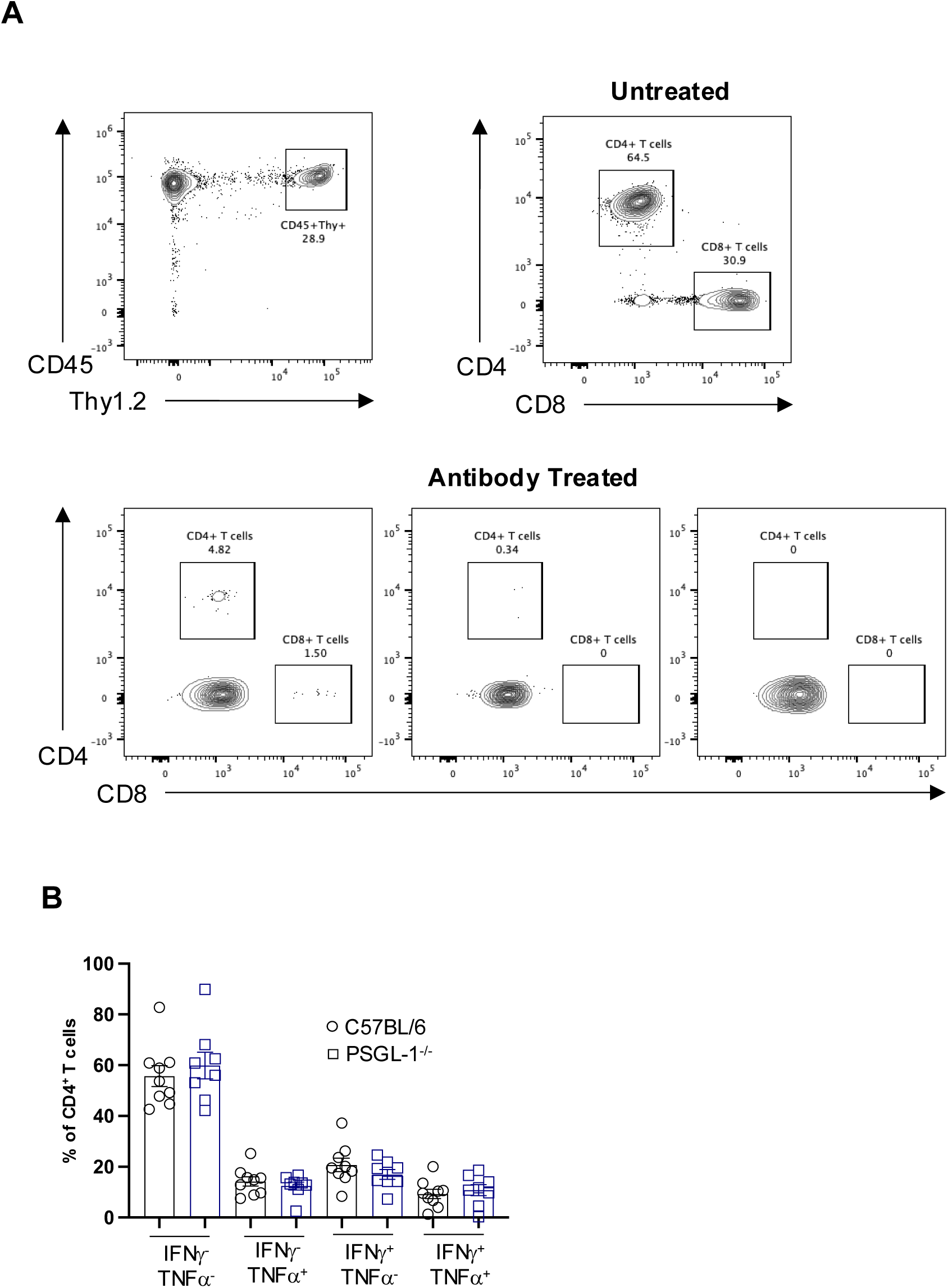
Control of KPC.4662 tumors in PSGL-1^-/-^ mice is T cell-dependent. **(A)** Representative FACS plots confirming CD4^+^ and CD8^+^ T cell depletion in three representative mice after receiving antibody treatment on days -2 and -1 prior to tumor inoculation. **(B)** Dot plot/bar graph of cytokine-producing intratumoral CD4^+^ T cells from C57BL/6 or PSGL-1^-/-^ mice after 4 hours of stimulation with PMA/ionomycin in the presence of Golgi inhibitors.

